# Alzheimer’s disease: the large gene instability hypothesis

**DOI:** 10.1101/189712

**Authors:** Sourena Soheili-Nezhad

## Abstract

All drug trials of the Alzheimer’s disease (AD) have failed to slow the progression of dementia in phase III studies, and the most effective therapeutic strategy remains controversial due to the poorly understood disease mechanisms. For AD drug design, amyloid beta (Aβ) and its cascade have been the primary focus since decades ago, but mounting evidence indicates that the underpinning molecular pathways of AD are more complex than the classical reductionist models.

Several genome-wide association studies (GWAS) have recently shed light on dark aspects of AD from a hypothesis-free perspective. Here, I use this novel insight to suggest that the amyloid cascade hypothesis may be a wrong model for AD therapeutic design. I review 23 novel genetic risk loci and show that, as a common theme, they code for receptor proteins and signal transducers of cell adhesion pathways, with clear implications in synaptic development, maintenance, and function. Contrary to the Aβ-based interpretation, but further reinforcing the unbiased genome-wide insight, the classical hallmark genes of AD including the amyloid precursor protein (APP), presenilins (PSEN), and APOE also take part in similar pathways of growth cone adhesion and contact-guidance during brain development. On this basis, I propose that a disrupted synaptic adhesion signaling nexus, rather than a protein aggregation process, may be the central point of convergence in AD mechanisms. By an exploratory bioinformatics analysis, I show that synaptic adhesion proteins are encoded by largest known human genes, and these extremely large genes may be vulnerable to DNA damage accumulation in aging due to their mutational fragility. As a prototypic example and an immediately testable hypothesis based on this argument, I suggest that mutational instability of the large Lrp1b tumor suppressor gene may be the primary etiological trigger for APOE/dab1 signaling disruption in late-onset AD.

In conclusion, the *large gene instability hypothesis* suggests that evolutionary forces of brain complexity have led to emergence of large and fragile synaptic genes, and these unstable genes are the bottleneck etiology of aging disorders including senile dementias. A paradigm shift is warranted in AD prevention and therapeutic design.

#### Glossary

AD: Alzheimer’s disease;
APP: Amyloid precursor protein;
GWAS: Genome-wide association study;
FAK: Focal adhesion kinase;
PSD: Postsynaptic density;
PSEN1/2: Presenilin1/2;
SFK: Src family kinase

## Introduction

More than a century has passed since the first report of a presenile dementia case by Alois Alzheimer^1^, and the current understanding of AD pathophysiology borrows from identification of the Aβ peptide as the main constituent of senile plaques and subsequent discovery of APP and PSEN mutations in rare familial forms of AD^2,3^. These observations were compiled to the amyloid cascade hypothesis in the pre-genomic era^4^, which remains the central theory of AD etiopathogenesis and implicates Aβ and neurofibrillary tangles as the causes of disease.

Nevertheless, due to methodological difficulties, Aβ species has hardly been validated as the causal force of neurodegeneration in humans. Despite the general support received from preclinical models, manipulating pathways of Aβ generation and clearance has yielded disappointing results in several clinical trials so far^5^. While a handful of clinical failures do not necessarily warrant disproval of a theory *per se*, overemphasis on a single disease model is a dangerous gamble and could be one of the many explanations for the lack of progress in AD therapeutic design^6^.

Accuracy of the amyloid cascade hypothesis is a topic of ongoing debate^7-12^, and the long-standing over-reliance on a potentially wrong model warrants development of independent mechanistic explanations for this prevalent cognitive disorder. For this aim, the novel genome-wide insight into AD risk loci provides a strong basis, since in contrast to the neuropathological hallmarks including senile plaques and neurofibrillary tangles, which are of questionable etiological significance^13^, genetic risk factors temporally precede earliest stages of brain development, aging, and degeneration, and are expected to inform on causal events in the disease cascade.

Genetic architecture of common late-onset AD is highly multifactorial and only partly understood. Although a number of susceptibility loci have been identified by genome-wide association studies^14-19^, mechanistic interpretation of these new observations have generally been under powerful influence of the amyloid cascade theory so far. In contrast, our report servers to provide an evidence-based framework for compiling the genetic pathways of AD within an Aβ-independent domain. The rest of this manuscript is organized as follows; in the first section, I aim to comprehensively revisit roles of classical and novel genetic modifiers of AD risk in pathways of normal cell physiology. I show that APP, presenilins and APOE as well as 23 other AD risk genes converge to common pathways of cell-extracellular adhesion signaling, with important implications in synaptic circuit formation and neurite outgrowth navigation. In the second section, I provide bioinformatics evidence for interaction of aging with this genetic landscape by showing that even the insidious “normal” rate of DNA damage in aging cells may disproportionately hamper synthesis of extremely large synaptic adhesion proteins in late life. Finally, several immediately testable predictions are provided for assessment of this new disease model.

### 1.1 The APP family genes encode evolutionarily-conserved cell adhesion proteins

Derailed catabolism of the APP protein and generation of an aggregation-prone Aβ species abstract the mainstream theory of AD pathophysiology, and several efforts have been made to block this cascade by means of Aβ immunotherapies or design of secretase inhibitors^5^. In contrast, three decades after successful cloning of the APP gene^20^, the potential physiological roles of its protein product remain under-explored and unknown.

APP codes for a single-pass transmembrane protein and shows high expression levels at the site of neuronal growth cones, structures that form motile tips of the outgrowing axons and dendrites in the developing brain^21^. The Aβ peptide enhances interaction of neurites with extracellular adhesion molecules and promotes elongation of cell membrane projections^22,23^. The full-length and membrane-tethered form of the APP protein also interacts with the extra-cellular matrix adhesion molecules including laminin, heparan sulfate, fibronectin and collagen^24-26^. More specifically, interaction of APP with laminin^24^ and heparan sulfate^27^ has neurite-promoting effects, and this protein stimulates assembly of hippocampal connections^28^. On the other hand, antisense-downregulation of APP inhibits extension of neurites^29^. APP demonstrates a dose-effect in affecting growth cone adhesion and guidance^30^. Increased dosage of APP in Down syndrome results in emergence of faster advancing growth cones with promoted adhesive properties and larger sizes^31^. In contrast, knockdown of the APP gene in zebrafish results in neurite outgrowth disruption^32^. Intriguingly, although wild-type human APP can rescue this abnormal phenotype, the mutated APP gene of familial AD fails to substitute for the normal function of animal gene^32^.

Several intracellular pathways are speculated to mediate the neurite-promoting effects of APP in neuronal membrane. The netrin pathway of neurite guidance incorporates APP as a co-receptor for cell signaling^33^. In this context, APP inactivation disrupts normal netrin signaling and diminishes axonal outgrowth^34^. APP also binds the extracellular reelin glycoprotein, which is a large adhesion molecule for guidance and migration of neurons^35^. Interaction of reelin with APP promotes outgrowth of hippocampal neurites^35^, and this functional interaction requires presence of another cell adhesion molecule, the α3β1-integrin, as well^35^. Of note, integrin receptors are the main component of focal adhesion complexes, and they co-localize with the APP protein^36,37^ at dynamic neuronal adhesion sites^38^. In line, interaction of integrin with APP modulates neuritic outgrowth^39^. Integrin also acts as an accessory reelin receptor for cell adhesion modulation and neuronal migration^40-42^, and therefore they functionally link two important AD risk genes, including APP and the APOE receptor pathway as shall be discussed later.

In addition to influencing growth cone movement, the APP protein also coordinates spatial migration of neurons during brain development^43^. Triple-knockout of the APP family genes in mice results in a neuronal migration defect similar to human lissencephaly^44^. Further implicating a potential role in cell migration, two candidate extracellular ligands of the transmembrane APP protein including pancortin and lingo1 orchestrate migration of neural precursor cells^45-47^. It is noteworthy that pathways of growth cone adhesion and cell movement are mechanistically convergent, since both of these biological motility events rely on specialized membrane protrusions, namely filopodia and lamellipodia, for changing extracellular adhesion forces and cell membrane reshaping. These membrane projections possess surface adhesion receptors, which control dynamic rearrangement of the intracellular actin cytoskeleton for changing cell polarity, shape and movement direction^48^.

In close homology to canonical pathways of cell adhesion, mounting evidence indicates that the cytoskeletal system is an important point of convergence in the APP signaling axis. Transmembrane APP is selectively localized to the cytoskeletal-rich regions of neuronal growth cones at dynamic adhesion sites^38,49^, and the APP intracellular domain (AICD) reportedly affects rearrangement of the cellular actin cytoskeleton^50^. In this context, AICD interacts with a number of intracellular signal transducers, including Fe65, Tip60, KAI1, DISC1, dab1, X11, and Grb2^51-53^. All of these signal transducers influence pathways of cytoskeletal rearrangement and cell movement in diverse cellular mechanisms spanning cancer cell migration and brain development:

- Fe65 and Tip60 affect the cytoskeletal system and moderate cancer cell migration^54^.
- KAI1 suppresses cancer cell migration by influencing cytoskeletal assembly^55,56^.
- DISC1 coordinates remodeling of the actin cytoskeleton in migrating neurons and growth cone-like protrusions^57^. Of note, this protein rescues neuronal migration defects caused by loss of the APP gene^51^.
- Dab1 is a mandatory adaptor of the lipoprotein receptors axis in the APOE/reelin signaling pathway and controls remodeling of the actin cytoskeleton in neuronal migration^58^.
- X11 is a recently discovered modulator of the reelin pathway and affects cell movement^59^.
- Grb2 is an adaptor molecule which links various receptors including integrins with intracellular pathways of cytoskeletal plasticity, and thereby regulates cancer cell migration^60,61^.

In line, there is also ample evidence for functional engagement of the APP family proteins in migration and invasion of various cancer cells through the cytoskeletal pathway^62,63^. Through a feedback-like mechanism, the cytoskeletal regulator Rac1 controls expression of the APP gene in primary hippocampal neurons^64^. This functional engagement in cell migration has probably been evolutionarily conserved, as the APP gene paralogue of Drosophila (APPL) has promoted the neuronal migration process since the earliest stages of nervous system evolution^65^. In line, phylogenetic evolution suggests that cell adhesion is the most consistent biological function of the APP family genes^66^.

The cytoplasmic tail of APP is noteworthy in the evolutionary context, since it comprises a super-conserved NPxY amino acid motif in the form of _682_YENPTY_687_ which has remained unchanged from roundworms to humans for more than 900 million years^67^. This consensus motif is known to mediate endocytic sorting of membrane receptors and their interaction with intracellular tyrosine-phosphorylated signaling adaptors^68^. Two mentioned intracellular adaptors of the APP protein, including dab1 and Fe65, interact with this consensus motif in a phosphorylation-dependent manner^69,70^. Further implicating a signaling role, the _682_Tyr residue of this APP motif undergoes phosphorylation and is essential to synaptogenesis^71^.

In addition to neurodevelopmental roles, the APP protein is also evidenced to maintain its function in mature neurons. Mouse hippocampal neurons express the APP protein under physiological conditions^72^, and APP is present in close proximity to post-synaptic NMDA glutamate receptors. APP controls postsynaptic trafficking of these synaptic receptors and promotes neurotransmission^73,74^. Through its conserved NPxY motif, APP also interacts with the postsynaptic scaffold protein AIDA-1^75^, which is a regulator of synaptic transmission and palsticity^76^. On the other hand, loss of the APP family genes disrupts synaptic function^77^, memory formation^78^, and causes an aging-related synaptic loss in mice^79,80^. APP and the other two members of this protein family form trans-synaptic adhesion dimers^81^. Cleavage of the APP protein changes synaptic adhesion and assembly^82^, and mutations in APP disrupt synaptic adhesion^83^. A more detailed review of the APP protein and its roles in neurophysiology is beyond the scope of this manuscript and the interested reader is referred to recent publications^84-86^.

### 1.2 The γ-secretase complex is a membrane-tethered enzyme for signaling of cell adhesion receptors

PSEN1 and PSEN2 genes code for catalytic subunits of the transmembrane γ-secretase enzyme, and various mutations in these genes underpin autosomal-dominant forms of AD. As a mandatory step in Aβ_40/42_ generation, γ-secretase cleaves the APP protein at the γ-site. However, as a surprising finding, it was recently observed that some PSEN mutations of familial AD cause an almost complete loss of γ-secretase function^87^ and reduce generation of the putatively-neurotoxic Aβ_40_, Aβ_42_ and Aβ_43_ species occasionally to undetectable levels^88,89^. In further contradiction, when knock-in mouse models were constructed using the mutated PSEN1 gene of familial AD, they were phenotypically similar to knockout strains lacking any γ-secretase function, with both of these strains demonstrating impaired hippocampal plasticity^90^. This novel line of evidence reinforces a loss-of-function impact for the PSEN mutations of familial AD, and may explain the paradoxical worsening of cognitive function and accelerated brain atrophy in the γ-secretase inhibitor trials of AD^91,92^.

In contrast to the narrow focus on derailed pathways of APP catabolism, unbiased proteomic profiling reveals that the γ-secretase enzyme has a broad spectrum of substrate specificity to molecules with transmembrane signaling roles^93,94^. For instance, the γ-secretase cleaves the APOE/reelin receptors^95^, as well as DSG2, TREM2, ephrin, and notch3 receptors^96^, which are all coded by AD risk genes^93,97-99^. Moreover, loss of γ-secretase has functional implications in neurobiology, and results in erroneous axonal pathfinding due to impaired netrin signaling^100^. Importantly, loss γ-secretase also disrupts cell adhesion force generation^101^.

Recent nanoscale microscopy has revealed that expression of the γ-secretase enzyme is selectively enriched in postsynaptic sites during normal synaptic maturation^102^. A synaptic role for the γ-secretase complex is further supported by its functional interaction with the glutamate receptors, as well as σ-catenin and N-cadherin which are synaptic adhesion molecules^102,103^. In this context, cleavage of cell adhesion receptors by the γ-secretase modulates synaptic adhesion and neurotransmission^103^. Familial AD mutations of presenilin disrupt this modulatory effect^104^.

### 1.3 The APOE-lipoprotein receptor axis coordinates contact-guidance of neuronal growth cones

APOE4 is the strongest genetic risk factor of common late-onset AD, explaining ∼6% of the disease risk^105^. In contrast, the only correlation of the APP locus with late-onset AD has been recently reported in an Icelandic cohort, showing that a rare protective variant explains less than 0.6% of the disease risk at a sub-genome wide statistical level^106^, albeit this variant does not contribute to protecting from AD in the North American population due to the extremely low allele frequency^107^. Despite this highly disproportionate level of evidence, mechanistic interpretation of the strong APOE4 risk factor still mostly borrows from potential influences on pathways of Aβ clearance.

The APOE molecule binds to the family of lipoprotein receptors and thereby moderates cellular uptake of lipoprotein particles in various organs. However, lipoprotein receptors are not simple cargo transporters, and stimulate a comprehensive nexus of intracellular second messenger signals^108^. For instance, the two lipoprotein receptors of the reelin pathway are shared with APOE, including APOEr2 and VLDLr receptors. Activation of these receptors by reelin triggers phosphorylation of the intracellular dab1 adaptor, which binds to the consensus NPxY motif of the receptor intracellular domain^109^. Through dab1 activation, the reelin pathway affects various aspects of cell physiology, among which cytoskeletal remodeling and neuronal migration are central^110^. Importantly, the reelin pathway guides extension of hippocampal neurites^111^ and coordinates outgrowth of the perforant path which forms the major input fibers to the hippocampal formation^112^.

The APOE molecule shares its lipoprotein signaling receptors with reelin^113^, and mounting evidence indicates that APOE also undertakes a similar role in guiding outgrowth of developing neurites^113-117^. The neurite promoting effect of APOE is isoform-dependent, with the APOE3 isoform being a more potent neurite outgrowth inducer than the APOE4 risk isoform^115,117^.

Unlike reelin, the intracellular signaling pathway of the APOE molecule has been less investigated in neurons, but partly studied in other cells. In macrophages, APOE activates transducers of the reelin pathway including dab1 and PI3K^118^. In vascular pericytes, APOE affects rearrangement of the actin cytoskeleton and its knockdown deranges normal cell migration^119^. The APOE4 isoform also affects the proteomic signature of cytoskeletal regulators in peripheral nerves^120^. Taken together, this body of evidence suggests that the APOE molecule may signal through a reelin-like network by incorporating lipoprotein receptors and the cytoskeletal system for inducing cell adhesion and movement.

In addition to the strong association of the APOE locus with AD, other risk loci further reinforce relevance of lipoprotein receptors and their signaling path in this disease. Variants within the reelin gene are the top genetic correlate of AD-type neuropathology in postmortem human brains^121^. F-spondin (Spon1), which codes for a reelin domain-containing cell adhesion molecule, is correlated with the rate of cognitive decline in AD and also affects white matter microstructure in healthy humans^122,123^. Moreover, F-spondin interacts with the APP protein^124^, and this interaction serves to activate signaling of the reelin adaptor dab1 in ganglion cells^125^. Two new AD risk loci including Sorl1 and CLU respectively code for a lipoprotein receptor and a lipoprotein receptor ligand^126,127^. Sorl1 regulates cell migration^127,128^ and CLU activates various transducers of the reelin pathway including dab1 and PI3K/Akt in neurons^129^.

Apparently unrelated to their roles in lipid metabolism, lipoprotein receptors interact with the major postsynaptic scaffold protein PSD95 and take part in synaptic architecture^130-132^. Expression of the lipoprotein receptors affects synaptic density in hippocampal and cortical neurons^133^. Moreover, lipoprotein and neurotransmitter receptors interact with each other^130,132^ and activation of the lipoprotein receptor pathway by reelin promotes synaptic plasticity^134-136^. Specifically, a recent study shows that postsynaptic activity of APOEr2 is critical for dab1 phosphorylation and insertion of AMPA glutamate receptors at postsynapse for long-term potentiation^137^. Lipoprotein receptors also share several intercellular signal transducers with the APP protein, including X11, dab1, and Fe65^133,138^, potentially reflecting convergent signaling pathways.

### 1.4 AD susceptibility loci strongly implicate cell adhesion pathways

Familial early-onset AD which is caused by APP or PSEN mutations constitutes less than one percent of diagnosed patients. In contrast, several genome-wide association studies have recently revealed the complex polygenic landscape of common late-onset AD^14-19^. Remarkably, the majority of late-onset AD risk genes engage in pathways of cell adhesion, migration and contact-guidance:

- **DSG2** (Desmoglein-2, rs8093731) is a component of desmosomal cell adhesion complexes. DSG2 gene product interacts with β8-integrin and serves focal adhesion roles in endothelial cells and regulates cytoskeletal assembly^139^. DSG2 also controls cell motility, and its depletion affects migration of malignant melanoma cells^140^.
- **EPHA1** (rs11771145) codes for a member of the ephrin-A receptor family of neurite adhesion and guidance. EPHA1 moderates cell migration through integrin-linked kinase and the cytoskeletal remodeling pathway^141,142^. EPHA1 also affects invasion and metastasis of colorectal cancer cells^143^.
- **FRMD4A**^144^ and **FERMT2** (Kindlin-2, rs17125944) code for two members of the FERM domain family, which link integrin and focal adhesion kinase (FAK) with the intracellular actin cytoskeleton^145,146^. FERMT2 transduces cell adhesion signals and is engaged in malignant cell invasion^147^.
- **GAB2** (rs2373115), one of the earliest AD susceptibility loci to be discovered by genome-wide scan^14,148^, encodes a scaffolding protein acting downstream to the integrin signaling pathway. GAB2 regulates adhesion and migration of hematopoietic cells^149^ and also controls cytoskeletal remodeling in migrating breast cancer cells^150^.
- **CASS4** (Hepl, rs7274581) controls focal cell adhesion^151^ and the CAS family members take part in axon guidance by interacting with integrin^152^. CASS4 also affects cytoskeletal reorganization and moderates cancer cell invasion^151,153^.
- **CD2AP** (rs10948363) codes for an actin cytoskeleton binding protein^154^. CD2AP regulates focal adhesion of kidney podocytes at contact sites by linking membrane adhesion complexes with the intracellular actin cytoskeleton^155^.
- **PTK2B** (Pyk2, rs28834970) is a focal adhesion signal transducer and affects cytoskeletal remodeling^156,157^. PTK2B coordinates integrin-dependent migration of T-cells^158^ and promotes invasion of malignant glioma cells^159^.
- **PICALM** (rs10792832) is a clathrin adaptor protein and engages in membrane receptor trafficking^160^. Clathrin regulates endocytosis of synaptic vesicles and moderates trafficking of the glutamate receptors^161^. Unbiased gene-gene interaction analysis has revealed that the PICALM locus interacts with DOCK1 in AD^162^, which is an actin cytoskeleton regulator and affects cell movement^163^.
- **INPP5D** (SHIP-1, rs35349669) is a key modulator of the PI3K pathway. This protein regulates platelet adhesion by affecting integrin signaling^164^. INPP5D also coordinates movement of neutrophils in response to focal contact and adhesion^165^.
- **NYAP1** (rs1476679) codes for a signal transducer of the PI3K pathway. NYAP1 acts downstream to signaling of the contactin5 synaptic adhesion molecule and controls cytoskeletal remodeling in outgrowing neurites^166^. Of note, contactin5 also binds the amyloid precursor-like protein 1^167^.
- **Amphysin II** (BIN1, rs6733839) codes for a protein which binds to the cytoplasmic tail of integrin^168^ and neuronal focal adhesion kinase^169^ and is therefore probably involved in integrin-dependent cell adhesion. Moreover, Amphysin I, which has a high level of sequence similarity (71%) with this gene product, regulates outgrowth of hippocampal neurites^170^ and links endocytosis mechanisms to pathways of cytoskeletal remodeling^171^.
- **UNC5C**^172^ (rs137875858) codes for a receptor of the netrin pathway of axon guidance^173^. The netrin pathway incorporates α3β1-integrin and the Down Syndrome Cell Adhesion Molecule (DSCAM) in neuronal migration process and neurite outgrowth, respectively^174,175^.
- **TPBG**, a recently discovered AD risk gene^19^, modulates cell adhesion and movement^176,177^. TPBG localizes at focal adhesion sites in kidney podocytes and affects formation of actin stress fibers for cell remodeling^178^. Deletion of TPBG disrupts cadherin-dependent cell adhesion and suppresses cell migration^179^.
- **HBEGF**^19^ (rs11168036) encodes a protein which promotes integrin-dependent cell adhesion^180^. HBEGF also regulates focal adhesion kinase and by rearranging the actin cytoskeleton moderates cell migration^181^.
- **USP6NL**^19^ (RNTRE, rs7920721) modulates the integrin signaling axis and controls focal adhesion turnover, thereby acting as a “brake” in cell migration^182^.
- **TREM2** (rs75932628), a novel AD risk locus^183^, is known to interact with the plexin-A1 adhesion molecule^184^, which is an axon guidance receptor. Interaction of plexin-A1 with the TREM family has been suggested to moderate cell adhesion and movement through the cytoskeletal pathway^185^. The Plexin pathway also antagonizes the integrin signaling axis and inhibits cell movement^186^.
- **TTC3**, a novel familial late-onset AD locus, maps to the Down syndrome critical region^187^. TTC3 modulates β1-integrin signaling in malignant cells^188^ and its increased levels affects assembly of the actin cytoskeleton and thereby disrupts neurite extension^189^.
- **PLCG2**^190^ (rs72824905) codes for a phospholipase and is activated by integrin for cell migration^191^. Activation of PLCG2 downstream to the integrin pathway moderates adhesion of leukocytes^192^.
- **ABI3**^190^ (rs616338) affects the cytoskeletal pathway and participates in formation of membrane protrusions for cell motility^193^. Its binding partner, the ABI3 binding protein, interacts with integrin at focal adhesion sites and suppresses malignant cell migration^194,195^.

Taken together, the genetic architecture of AD strongly implicates various cell adhesion regulators and pathways of cytoskeletal plasticity. Further aiding in formulation of a unified disease model, many of these gene products cross-talk with the integrin pathway of focal adhesion. This convergence also strongly spotlights the Aβ-independent roles of the APP protein, γ-secretase and the APOE receptors in cell adhesion regulation and synaptic function.

## 2 The hypothesis

By using the unbiased genetic architecture of AD, our model puts the cell adhesion process at the center of disease pathways. Focal adhesion regulators including integrins coordinate cell migration, neurite outgrowth, and assembly of synaptic circuits in brain development. In the post-developmental brain, these canonical pathways also undertake pivotal roles in maintaining synaptic adhesion and plasticity^196^. Synaptic adhesion molecules form a dense scaffold at the postsynaptic density (PSD) sites and dendritic spines. This scaffold connects neurotransmitter receptors and ion channels with the intracellular actin cytoskeleton as well as the extracellular matrix, aiding in synaptic maintenance and dynamic remodeling.

Synaptic adhesion molecules also act as mechano-chemical sensors and actively moderate trafficking of neurotransmitter receptors^197^. For instance, it has been shown that enhancing signaling of the synaptic integrin receptors by application of an agonist peptide modulates neurotransmission^198^ in a dose-dependent manner^199^. In this context, integrin affects rearrangement of the actin cytoskeleton and promotes budding of filopodia-structures that strengthen synaptic connections^200^. Remarkably, this is the same mechanism through which the integrin pathway coordinates growth-cone adhesion and pathfinding during synaptic circuit development^201^. It is noteworthy that the post-developmental role of cell adhesion pathways in synaptic physiology is not limited to integrins, and has been observed for several cell adhesion molecules (Fig. 1).

**Figure 1.**
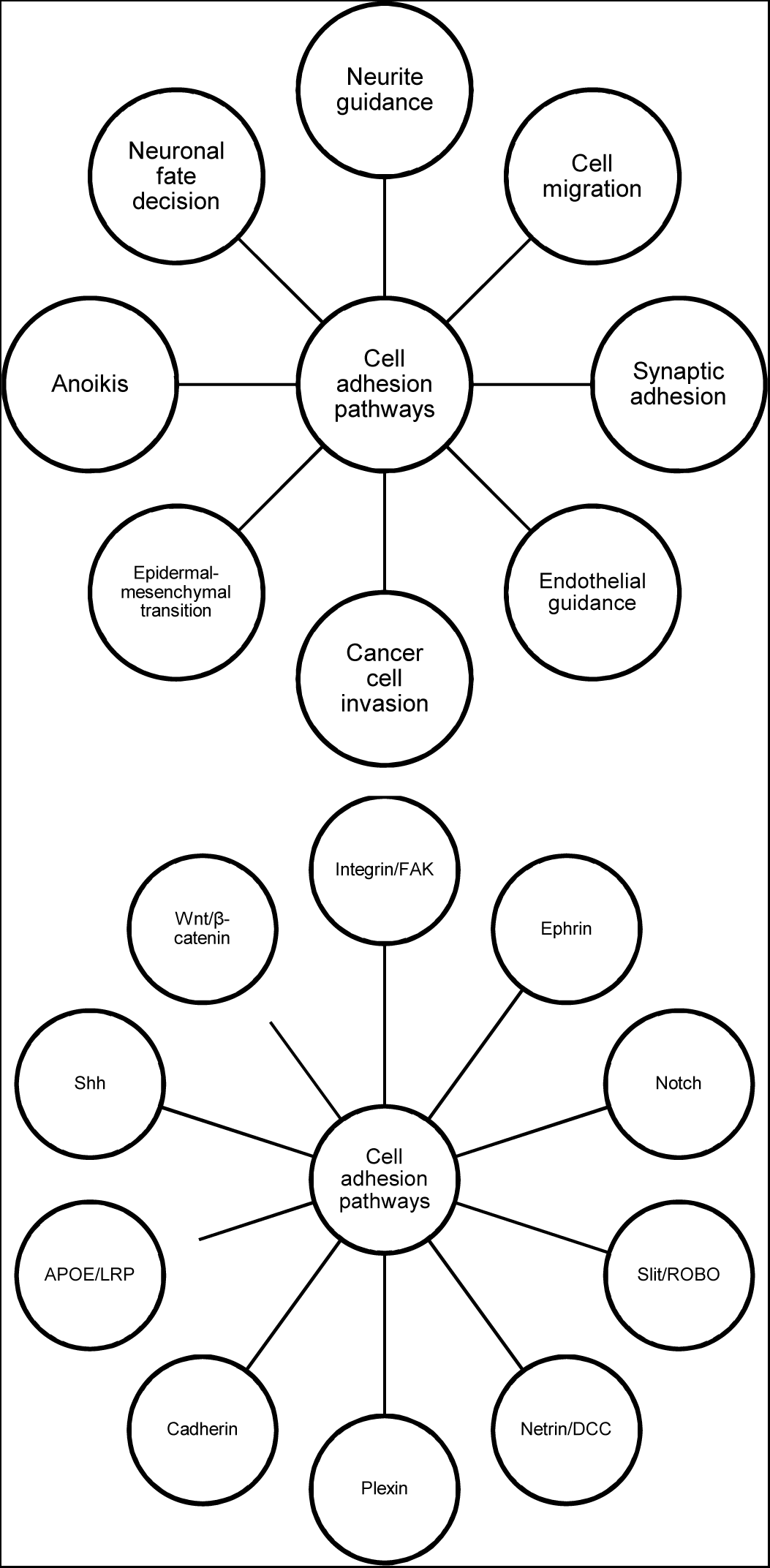
Biological adhesion pathways transfer extracellular signals across the cell membrane, and affect cell polarity, movement and survival (top). Various pathways of extracellular adhesion signaling coordinate rearrangement of the actin cytoskeleton and thereby control reshaping of membrane projections for cell movement and plasticity (bottom). FAK: focal adhesion kinase; LRP: lipoprotein receptor; Shh: Sonic hedgehog.

I propose that the heritable component of AD is determined by genetic factors which coordinate growth cone adhesion and assembly of synaptic circuits in brain development. The same molecular machinery also takes part in post-developmental synaptic maintenance, plasticity and functional resilience in later life. In this regard, any factor causing disruption of biological adhesion pathways in aging may lead to synaptic failure and cognitive decline.

## 3 Aging and Alzheimer’s disease

Human aging is the strongest risk factor for various dementias including AD. Considering the high prevalence of AD in late life, this disease may represent a continuation of global aging process, and cellular disruptions which happen in “normal” aging may give rise to AD when accelerated^7^. An elegant work has recently revealed that frontal cortex cells of healthy humans accumulate ∼37 new point mutations each year^202^, and these mutations may represent the final outcome of a broader DNA damage process. Loss of genomic integrity is one of the factors already implicated in AD etiopathogenesis, but its relevance to molecular disease pathways has not been elucidated^12,203,204^.

From a statistical point of view, even if a fully random process causes accumulation of mutations in aging neurons, larger genes are expected to be disproportionately affected in late life. Suppose that the burden of 37 annual mutations is uniformly scattered at purely random genomic positions in neurons (5.7 ×10^−9^ mutations/base pair.year). In this scenario, approximately 1% copies of a median-sized human gene (29.6kbp) will acquire at least one somatic mutation in a 65-year individual. In sharp contrast, the largest known human gene, CNTNAP2, which codes for a synaptic adhesion protein and is more than 80 × larger than the median-sized gene, is expected to be highly vulnerable to somatic mutations, and only 42% of its copies are estimated to remain intact in the same individual (Fig. 2). This high variability in the risk of mutations is due to the statistical distribution of gene sizes, which spans three-orders of magnitude with a long tail encompassing extremely large genes (Fig. 3).

**Figure 3.**
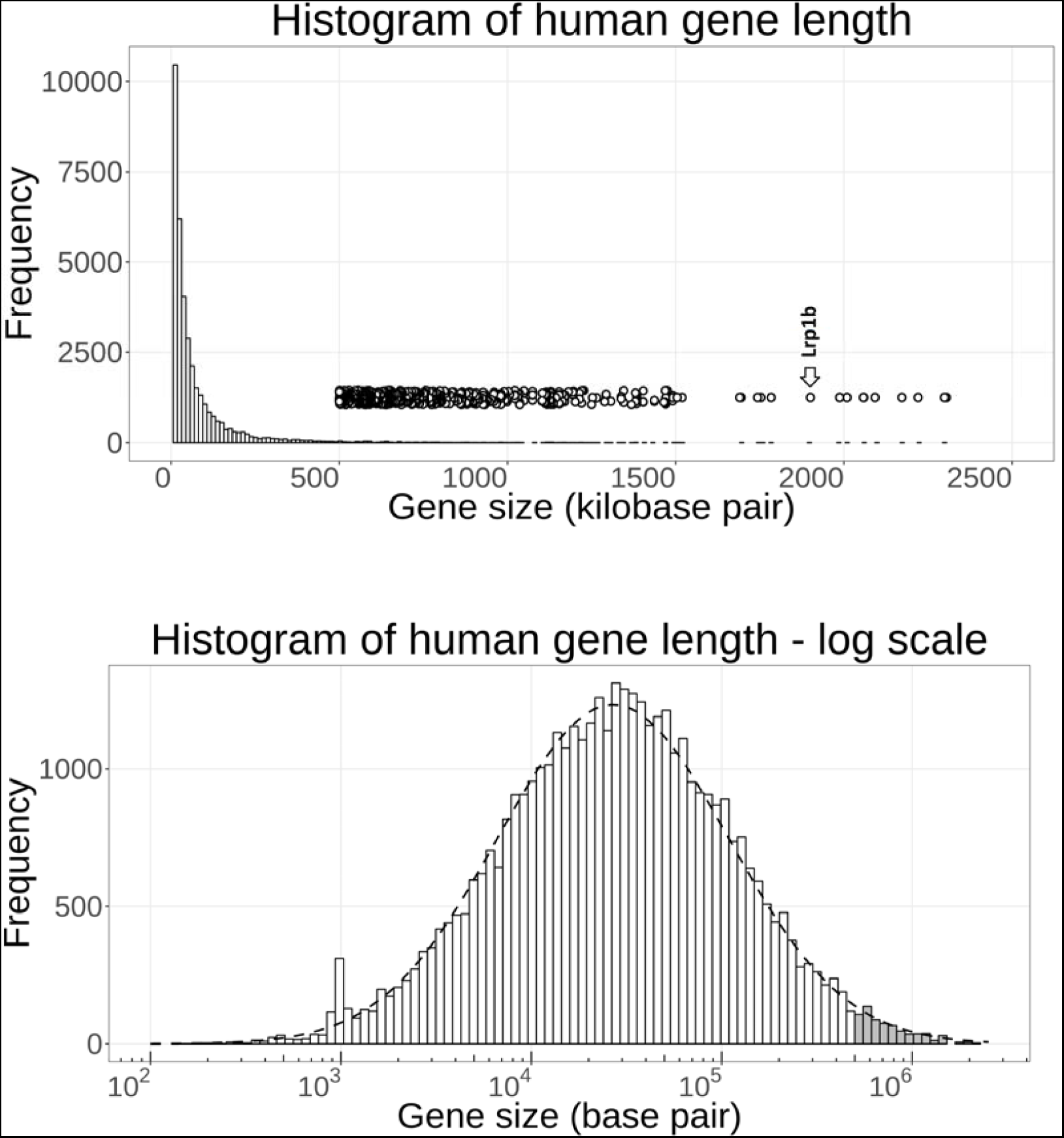
Human gene length distribution has a long tail which extends towards a group of extremely-large genes in the megabase pair range (top). The arrow points to the giant APOE receptor, Lrp1b. Human gene size parameter closely follows a log-normal distribution (bottom) with parameters μ=ln(26.9kbp) and σ=1.4. The outlier bin near 1 kbp represents the large family of olfactory receptors that have gone through extreme evolutionary expansion. Scattered circles (top) and grey bars (bottom) show the subgroup of large genes used in functional enrichment analyses of this paper (>500kbp).

**Figure 2.**
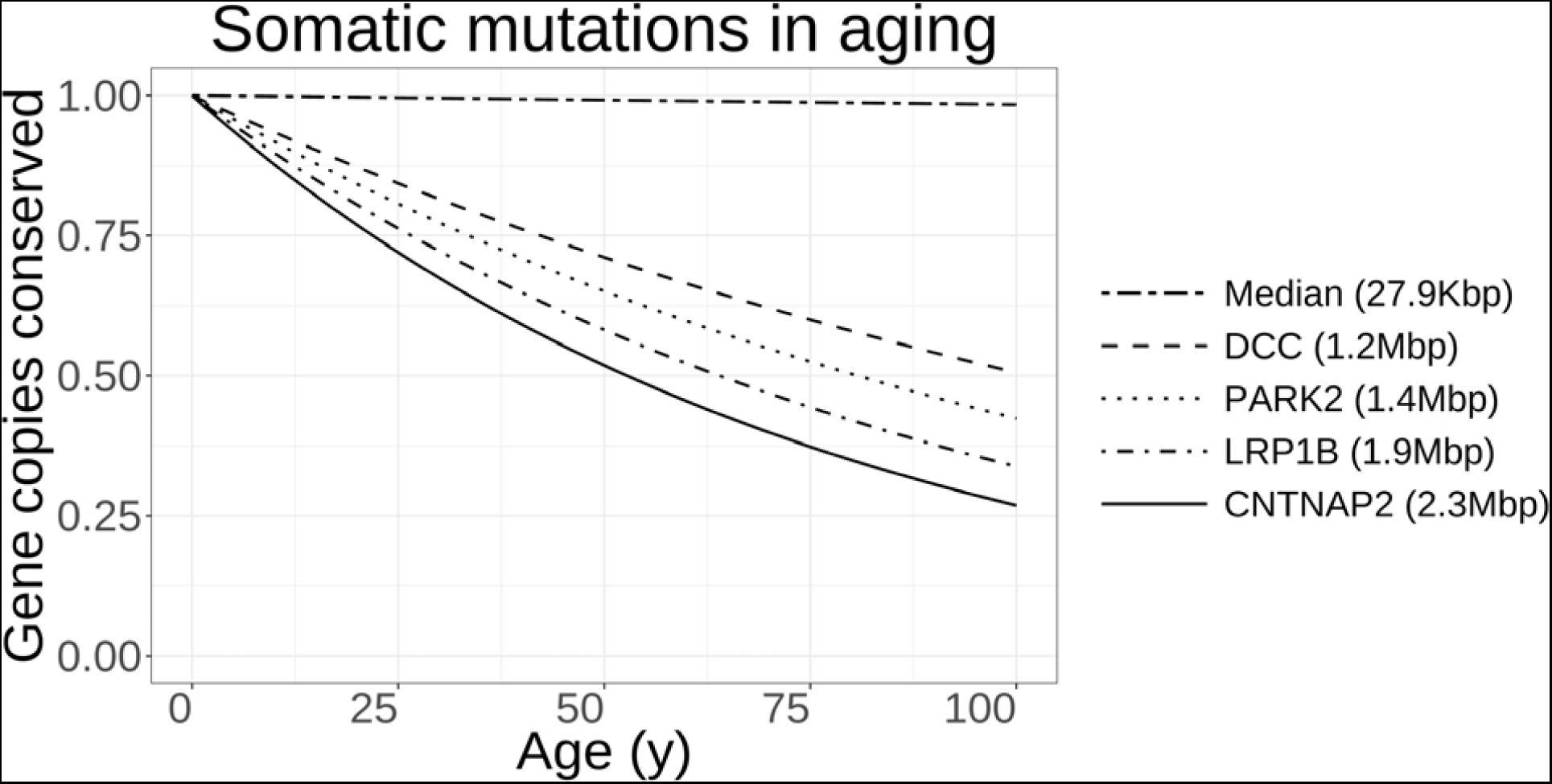
A simple binomial model in which somatic mutations take place at a fixed and uniform rate across the genome reveals that a median-sized human gene mostly survives the mutational burden of aging, with only ∼1% of its copies being affected by any somatic mutation in a 65 year-old subject. However, larger genes will have a significantly shorter *half-lives* set at the 6^th^ and 7^th^ decade of life; many of these large genes regulate synaptic adhesion and function with relevance to neurodegenerative disorders, and also act as fragile tumor suppressors.

Why has the evolution in some cases selected for extremely large genes, although they are known to map to chromosomal fragile sites^205^ and possibly be more vulnerable to DNA damage? I was compelled to objectively investigate whether large human genes non-randomly take part in certain biological themes, cellular functions, and tissue types for a potential explanation of their exceptional evolutionary trajectory. For this aim, I size-sorted all of the protein-coding human genes (n=19,287 RefSeq genes that successfully mapped to DAVID indices), and considered the gene length threshold of >500kb for defining *large* human genes. This cut-off threshold resulted in consideration of 260 large human genes representing 1.3% of all protein-coding transcripts. Functional annotation profile, pathway enrichment, and tissue expression of this gene set of interest were investigated using a standard DAVID query^206,207^.

Interestingly, the top overrepresented organs label for selective expression of these large genes were *brain* (p=1.4 ×10^−19^), followed by *amygdala* (p=3.1 ×10^−5^), and *hippocampus* (p=6.6 ×10^−5^). By showing strong enrichment statistics, *homophilic cell adhesion via plasma membrane adhesion molecules* was the most overrepresented biological process related to this gene set of interest (Table 1), and the most overrepresented cellular component was *postsynaptic membrane* (Table 2). All other enriched gene ontology terms further implicated pathways of nervous system development and physiology (Table 1). Among KEGG curated biological pathways, four pathways were found to be statistically enriched, including *Glutamatergic synapse* (hsa04724; corrected p=0.02), *Axon guidance* (hsa04360; corrected p=0.03), *Cell adhesion molecules* (hsa04514; corrected p=0.04), and *Insulin secretion* (hsa04911; corrected p=0.04).

**Table 2.** Enrichment of large human genes (>500 kbp) in *gene ontology: cellular component* annotations.

| Gene ontology term – cellular component | Gene count | p-value | Corrected p-value |
| --- | --- | --- | --- |
| GO:0045211~postsynaptic membrane | 22 | $2 \times 10^{-12}$ | $5 \times 10^{-10}$ |
| GO:0030054~cell junction | 29 | $7 \times 10^{-11}$ | $8 \times 10^{-9}$ |
| GO:0005886~plasma membrane | 100 | $3 \times 10^{-10}$ | $3 \times 10^{-8}$ |
| GO:0014069~postsynaptic density | 18 | $8 \times 10^{-10}$ | $5 \times 10^{-8}$ |
| GO:0042734~presynaptic membrane | 12 | $9 \times 10^{-10}$ | $4 \times 10^{-8}$ |
| GO:0030424~axon | 16 | $6 \times 10^{-7}$ | $2 \times 10^{-5}$ |
| GO:0045202~synapse | 14 | $2 \times 10^{-6}$ | $6 \times 10^{-5}$ |
| GO:0048786~presynaptic active zone | 7 | $3 \times 10^{-6}$ | $8 \times 10^{-5}$ |
| GO:0016021~integral component of membrane | 104 | $3 \times 10^{-6}$ | $7 \times 10^{-5}$ |
| GO:0043197~dendritic spine | 10 | $1 \times 10^{-5}$ | $3 \times 10^{-4}$ |
| GO:0031225~anchored component of membrane | 10 | $3 \times 10^{-5}$ | $6 \times 10^{-4}$ |
| GO:0043005~neuron projection | 14 | $3 \times 10^{-5}$ | $6 \times 10^{-4}$ |
| GO:0042383~sarcolemma | 8 | $2 \times 10^{-4}$ | $3 \times 10^{-3}$ |
| GO:0005856~cytoskeleton | 16 | $2 \times 10^{-4}$ | $4 \times 10^{-3}$ |
| GO:0030425~dendrite | 15 | $3 \times 10^{-4}$ | $4 \times 10^{-3}$ |

**Table 1.** Enrichment of large human genes (>500 kbp) in *gene ontology: biological process* annotations.

| Gene ontology term – biological process | Gene count | p-value | Corrected p-value |
| --- | --- | --- | --- |
| GO:0007156~homophilic cell adhesion via plasma membrane adhesion molecules | 19 | $1 \times 10^{-11}$ | $1 \times 10^{-8}$ |
| GO:0007157~heterophilic cell-cell adhesion via plasma membrane cell adhesion molecules | 11 | $2 \times 10^{-9}$ | $1 \times 10^{-6}$ |
| GO:0007155~cell adhesion | 27 | $2 \times 10^{-9}$ | $8 \times 10^{-7}$ |
| GO:0007399~nervous system development | 20 | $3 \times 10^{-8}$ | $9 \times 10^{-6}$ |
| GO:0007416~synapse assembly | 10 | $2 \times 10^{-7}$ | $5 \times 10^{-5}$ |
| GO:0007411~axon guidance | 14 | $4 \times 10^{-7}$ | $9 \times 10^{-5}$ |
| GO:0097120~receptor localization to synapse | 5 | $5 \times 10^{-6}$ | $9 \times 10^{-4}$ |
| GO:0007165~signal transduction | 37 | $6 \times 10^{-6}$ | $1 \times 10^{-3}$ |
| GO:0007612~learning | 8 | $2 \times 10^{-5}$ | $2 \times 10^{-3}$ |
| GO:0051965~positive regulation of synapse assembly | 8 | $3 \times 10^{-5}$ | $3 \times 10^{-3}$ |
| GO:0007158~neuron cell-cell adhesion | 5 | $6 \times 10^{-5}$ | $7 \times 10^{-3}$ |
| GO:0007269~neurotransmitter secretion | 7 | $8 \times 10^{-5}$ | $9 \times 10^{-3}$ |
| GO:2000463~positive regulation of excitatory postsynaptic potential | 5 | $2 \times 10^{-4}$ | 0.01 |
| GO:0097105~presynaptic membrane assembly | 4 | $2 \times 10^{-4}$ | 0.02 |
| GO:0051966~regulation of synaptic transmission, glutamatergic | 5 | $3 \times 10^{-4}$ | 0.02 |
| GO:0007420~brain development | 11 | $4 \times 10^{-4}$ | 0.03 |
| GO:0030534~adult behavior | 5 | $5 \times 10^{-4}$ | 0.03 |
| GO:0051491~positive regulation of filopodium assembly | 5 | $5 \times 10^{-4}$ | 0.03 |
| GO:0035176~social behavior | 6 | $6 \times 10^{-4}$ | 0.04 |
| GO:0007268~chemical synaptic transmission | 12 | $6 \times 10^{-4}$ | 0.04 |
| GO:0035418~protein localization to synapse | 4 | $6 \times 10^{-4}$ | 0.04 |

The strong selectivity of large human genes to brain, synapse and cell adhesion process is an enlightening observation, and I suggest that it may reflect existence of specialized natural selection forces for driving complexity of cognitive function in the evolutionary trajectory of organisms; these exceptionally large genes may have fostered adhesion and assembly of complex synaptic circuits in brain evolution. However, while larger genes may have promoted brain complexity, as an evolutionary bottleneck, they may also be inherently costlier to be maintained in late life due to limited DNA repair mechanisms, and such large synaptic genes may put modern humans at a neurobiological disadvantage when the burden of DNA damage is accumulated during the longer lifespan of modern humans. Importantly, since the average human life expectancy passed the 40-year milestone only two centuries ago^208^, there has been very weak evolutionary force for correcting the dementia-causing genomic variations. Taken together, rapid increase of brain complexity in parallel with extension of life expectancy may have recently unmasked a DNA maintenance and repair bottleneck in modern humans, which eventually presents as AD and potentially some other forms of senile disorders.

## 4 Predictions

Due to a combination of heritable factors and environmental exposures, AD patients may suffer faster accumulation of DNA damage in their neuronal genomes. This argument may be testable by revealing correlations between the longitudinal trajectories of cognitive decline in aging humans and the burden of somatic mutations in neurons. More specifically, a number of synaptic adhesion genes may be exceptionally vulnerable to DNA damage in certain neuronal populations. As a prototypic example, I predict that mutational instability of the Lrp1b gene in amygdalar and hippocampal neurons may be increased in the typical “APOE-type” sporadic AD patients (Fig. 4):

**Figure 4.**
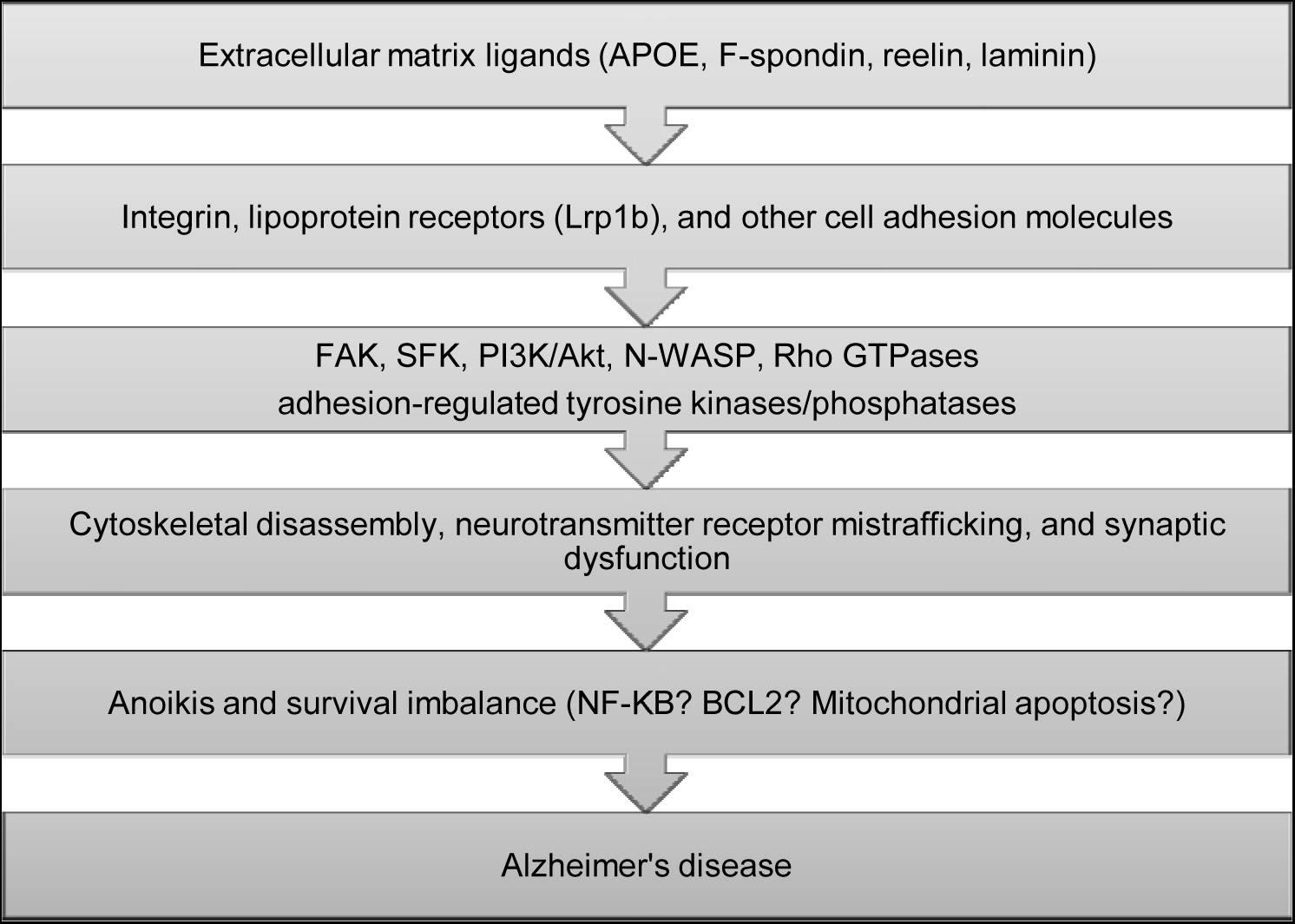
A simplified cascade of late-onset AD pathogenesis based on lipoprotein receptor signaling disruption. FAK: focal adhesion kinase; SFK: Src-family kinase.

- Lrp1b has affinity to both APOE and APP^209,210^.
- The Lrp1b gene demonstrates selective brain expression^209^ with hippocampal and amygdalar neurons showing the highest levels of Lrp1b transcription in humans^211^ (Fig. 5). Lrp1b also interacts with the major postsynaptic scaffold protein, PSD95^131^ as well as the synaptic plasticity-regulating protein PICK1^212^.

**Figure 5.**
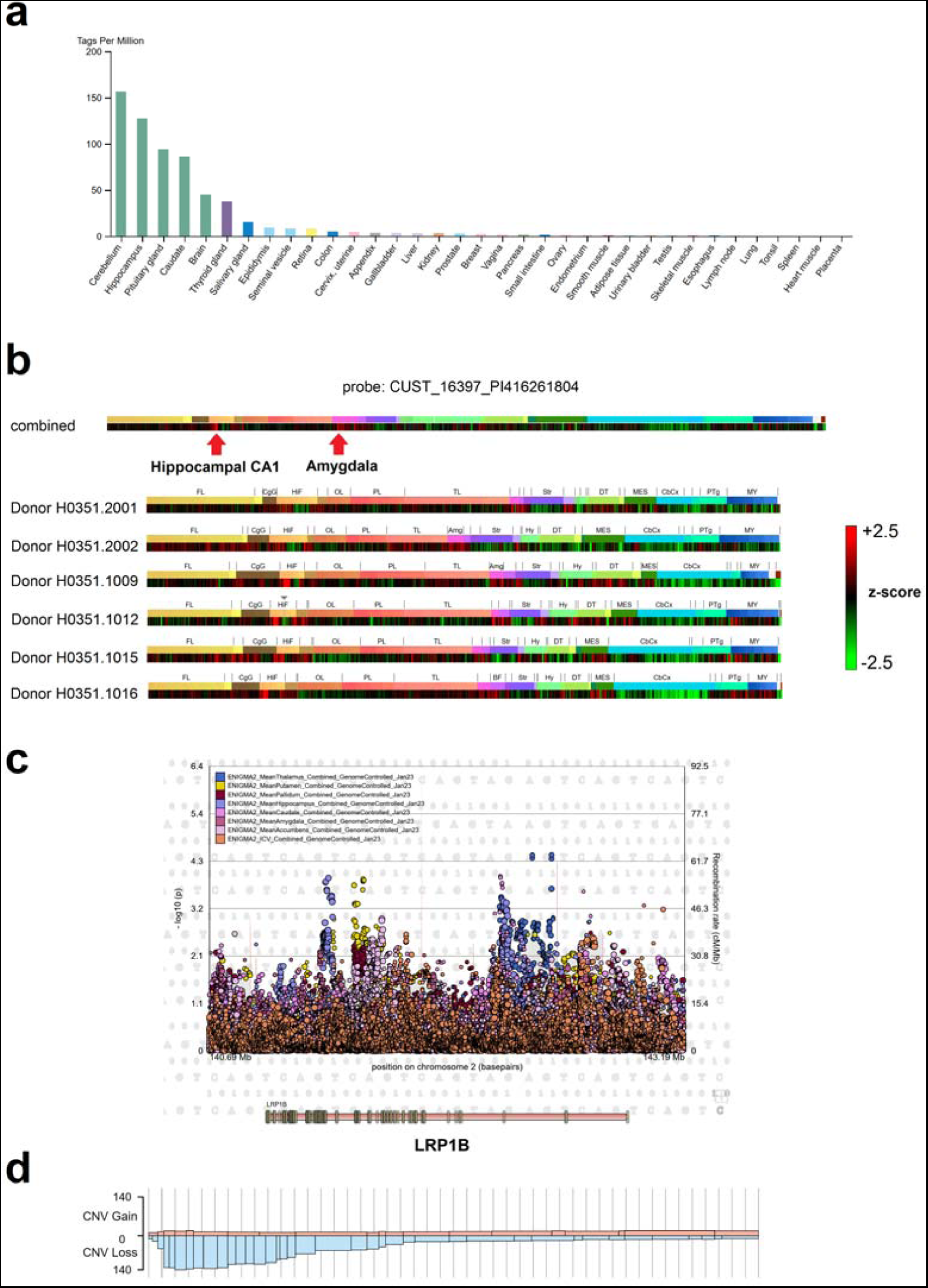
Tissue expression profile of Lrp1b in various organs (a) shows strong specificity to brain in FANTOM5 database. Spatial expression of Lrp1b in various brain structures in six postmortem human brain samples of the Allen human brain atlas (b). Correlation of genetic variants in the Lrp1b locus with several MRI measures of brain volume (c) in the ENIGMA-2 database^219,220^. Copy-number variation of the Lrp1b gene in 2,383 unique cancer tissue samples (d) shows a high probability of copy number loss in the 3` end of this gene.
- Lrp1b is the largest member of the lipoprotein receptor family genes and at an extreme size of 1.9Mbp is the 8^th^ largest human gene overall. Potentially due to its size and mapping to the chromosomal fragile site FRA2F, Lrp1b is among the ten most frequently deleted genes observed in a study of 3,131 cancer specimens^213^.
- The Lrp1b gene product controls focal adhesion, cytoskeletal remodeling and cell migration^214,215^, pathways which align with the genetic architecture of AD. Lrp1b is also cleaved by the γ-secretase enzyme and its intracellular fragment affects cell anchorage and survival^216^.
- Genetic variants of the Lrp1b locus are correlated with cognitive function in aging and AD^217,218^.

I predict that AD-type cognitive decline is correlated with propagation of DNA damage and somatic mutations in certain synaptic genes including Lrp1b, and subsequent dysfunctions in their intracellular pathways involving synaptic adhesion and maintenance. Although previous models have already implicated oxidative stress and DNA damage mechanisms in AD^12,203,204,221^, high-throughput results do not support an oxidative etiology for the observed mutations. Oxidative stress typically causes G:C→T:A transversions due to formation of free radicals^222,223^. However, aging cells demonstrate a clock-like signature of somatic mutations with enrichment of C:G→T:A transitions^224,225^. Intriguingly, this fingerprint was recently observed as the dominant type of mutations in neurons^223,226,227^. The reason for aging-related preponderance of C:G→T:A transitions is currently unknown, but spontaneous cytosine deamination, transcriptional stress, and failure of certain DNA repair mechanisms including base and nucleotide excision repair are potential explanations^228^.

It is noteworthy that Lrp1b only serves to provide one example of vulnerable synaptic genes in brain aging, and the true genetic landscape of AD and senile neurodegenerations is probably not reducible to the lipoprotein receptor axis (Fig. 6). Similar to loss of different tumor suppressor genes in various cancers which is caused by diverse DNA damage mechanisms, brain-wide expression of several unstable synaptic genes may underpin dementia heterogeneity in aging humans. For instance, the genome-wide landscape of the Parkinson’s disease implicates several genes of the synaptic vesicular trafficking system, including the extremely large tumor suppressor PARK2 mapping to the chromosomal fragile site FRA6E^229^. In this regard, dopaminergic neurons of substantia nigra are the most vulnerable structures in Parkinson’s disease, and they can be distinguished by selective expression of two tumor-suppressor genes with cell adhesion roles, including DCC^230^ and AJAP1^231^.

**Figure 6.**
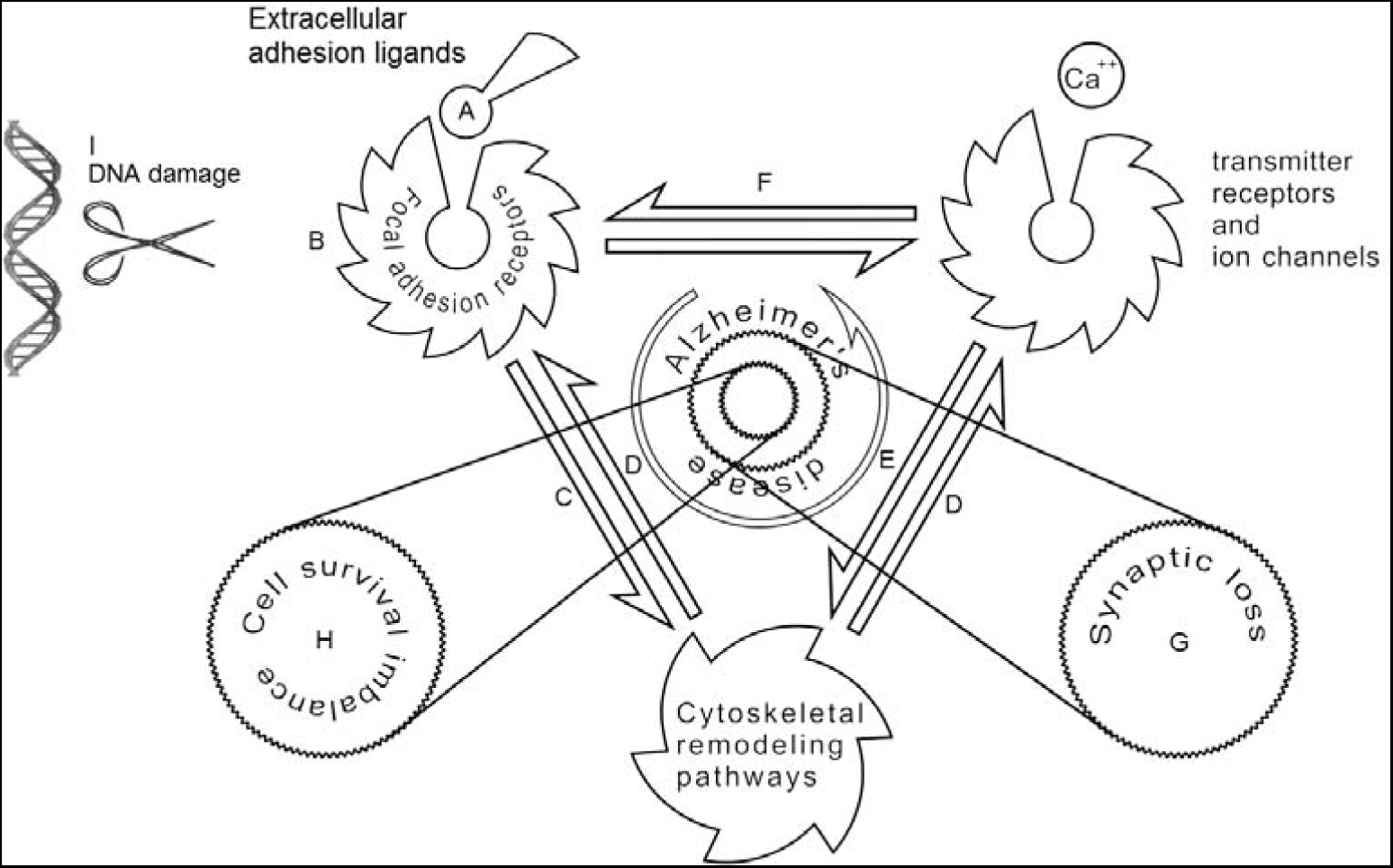
The proposed mechanisms of synaptic loss and neuronal death in AD. The extracellular matrix and cell adhesion molecules (A) modulate signaling of neuronal adhesion receptors (B). Cell adhesion pathways affect remodeling of synaptic actin cytoskeleton as well as other mediators of plasticity, e.g. various SH3 domain containing proteins (C). The postsynaptic density is anchored to the synaptic actin cytoskeleton through scaffolding proteins, e.g. PDZ domain containing proteins (D). Normal function and trafficking of the neurotransmitter receptors are controlled by cytoskeletal plasticity pathways (E) as well as membrane adhesion complexes (F). Disruption of cell adhesion pathways in AD impairs synaptic stability and causes dendritic spine loss (G), and may eventually lead to neuronal survival imbalance by triggering anoikis cascades (H). Selective vulnerability of genes with extremely large sizes or other features causing mutational instability may be the etiology of cell adhesion disruption in aging (I).

## 5 Future perspectives

Mice with distal truncation of Lrp1b have no apparent phenotype^131^, but a more proximally truncated Lrp1b causes early embryonic lethality^232^. Intriguingly, conditional knockout of the Lrp1 gene with 52% amino acid similarity to Lrp1b results in neurodegenerative changes in animals after 12 months of aging^233^. Conditional knockout of the Lrp1b gene and other modulators of the reelin/lipoprotein receptor signaling axis after completion of brain development may aid in modeling AD-type synaptic loss in animals.

Since even the most aggressive forms of AD remain clinically silent for decades, accelerating the aging process in laboratory animals may be necessary, for instance by crossing AD models with transcription-coupled DNA repair defective strains^234^ or usage of mutagenic forces such as UV radiation.

Our hypothesis is not based on any form of etiological relevance for Aβ species, amyloid plaques or neurofibrillary tangles in causal disease pathways, and redefines these pathological features as bystander epiphenomena. Even the strong APOE risk locus of sporadic AD fails to explain ∼94% of the disease variance. Therefore, single pathway therapeutic approaches may provide limited benefit in clinical trials.

In conclusion, this proposal, the *large gene instability hypothesis,* implicates DNA damage accumulation and loss of fragile synaptic adhesion genes as the primary etiology of AD. A shift of paradigm is warranted in AD drug design from manipulating the protein aggregation process to genetic engineering strategies such as large capacity gene therapy vectors.

